# Accurate RNA velocity estimation based on multibatch network reveals complex lineage in batch scRNA-seq data

**DOI:** 10.1101/2023.11.19.567699

**Authors:** Zhaoyang Huang, Xinyang Guo, Jie Qin, Lin Gao, Fen Ju, Chenguang Zhao, Liang Yu

**Author notes:** **Email:** (LY); (CZ).

## Abstract

RNA Velocity, as an extension of trajectory inference, is an effective method for understanding cell development using single-cell RNA sequencing (scRNA-seq) experiments. Nevertheless, existing RNA velocity methods are limited by the batch effect because they cannot directly correct for batch effects in the input data, which comprises spliced and unspliced matrices in a proportional relationship. This limitation can lead to incorrect velocity graphs. This paper introduces VeloVGI, which addresses this issue innovatively in two key ways. Firstly, it employs an optimal transport (OT) and mutual nearest neighbor (MNN) approach to construct neighbors in batch data. This strategy overcomes the limitations of existing methods that are affected by the batch effect. Secondly, VeloVGI improves upon VeloVI’s velocity estimation by incorporating the graph structure into the encoder for more effective feature extraction. The effectiveness of VeloVGI was demonstrated in various scenarios, including the mouse spinal cord and olfactory bulb, as well as on several public datasets. The results showed that VeloVGI outperformed other methods in terms of metric performance.

**Significance Statement:** RNA Velocity is an effective method for understanding cell development using single-cell RNA sequencing (scRNA-seq) experiments. This paper introduces VeloVGI, which addresses this batch effect issue for existing RNA velocity methods. The effectiveness of VeloVGI was demonstrated in various scenarios, including the mouse spinal cord and olfactory bulb, as well as on several public datasets. The results showed that VeloVGI outperformed other methods in terms of metric performance.

## Introduction

Single-cell RNA sequencing (scRNA-seq), a cutting-edge technology in the realm of single-cell genomics, enables the profiling of individual cells at the transcriptome level. Nevertheless, a significant hurdle in this field lies in capturing dynamic processes, such as cell type transitions, from static snapshots. Gaining insights into these transitions is pivotal for deciphering intricate phenomena like cell differentiation and cycle progression during development[1].

Numerous trajectory inference methods have been developed at the methodological level[2]. However, these methods have certain limitations as they solely describe the current snapshot and lack predictions of both past and future states. To address this, recent advancements have been made in trajectory inference using RNA velocity[3]. This approach leverages the dynamic changes from nascent to mature mRNA splicing, establishing a proportional relationship between the two to describe the dynamic trends within cells. By correlating cells and reflecting the differentiation relationships between them, RNA velocity provides insights into past and future states. Visualization of the results reveals that each cell possesses a vector, where the direction and length of the vector represent the direction and intensity of differentiation, respectively. Existing methods can be broadly categorized into two groups: machine learning and statistics: velocyto[3], scVelo[4], CellRank[5], Dynamo[6]. Deep Learning: VeloAE[7], UnitVelo[8], DeepVelo[9], VeloVAE[10], Pyro-velocity[11], LatentVelo[12]. These methods collectively contribute to our understanding of single-cell dynamics and play a crucial role in characterizing cell differentiation processes.

Current studies in single-cell RNA sequencing (scRNA-seq) analysis and cell atlases often involve collecting multiple samples across different experimental conditions and locations, aiming to shed light on a wider range of biological phenomena. Time-series sample analysis is effective for understanding cell differentiation[13], but it introduces batch effects. Existing RNA velocity methods can produce error-prone velocity graphs when batch effects are overlooked, even after preprocessing with batch correction techniques[14]. This issue arises because batch integration tools typically process a single expression matrix, while RNA velocity quantification tools[3, 15, 16] provide separate matrices for spliced and unspliced mRNA expressions. Correcting these matrices separately or concatenating them can disrupt the relative ratios of the two types of mRNA, leading to inaccurate results[17]. Therefore, there is a pressing need to develop RNA velocity methods specifically designed for multi-batch scRNA-seq datasets, which is the primary focus of this study.

In a recent study[18], it was demonstrated that the neighborhood construction process during preprocessing significantly impacts the final RNA velocity results. The presence of batch effects naturally leads to more cell-cell neighbor relationships within the same batch and fewer neighbor relationships between different batches in traditional KNN (K nearest neighbor) approaches. To address this issue, we drew inspiration from Waddington-OT[19] to compute optimal transportation for adjacent batches and used MNN (mutual nearest neighbor)[20] to establish inter-batch neighbor relationships. We manually specified these relationships based on the optimal transportation method and mutual nearest neighbor. Additionally, we incorporated the effectiveness of variational autoencoder (VAE) in removing batch effects [21]. Building upon this, we proposed VeloVGI which enhanced the encoder part of VeloVI[22], performing feature extraction on the fine-tuned graph structure to estimate RNA velocity for all batches. Furthermore, our method incorporates sampling and aggregation strategies, along with the inductive minibatch approach GraphSAGE[23], during model training to reduce computational overhead. We conducted a series of downstream analyses to demonstrate the effectiveness of our model results.

We conducted extensive tests on a variety of datasets to evaluate the performance of our approach. These datasets included the mouse spinal cord and olfactory bulb, as well as several publicly available datasets. Our method consistently demonstrated the ability to accurately capture distinct differentiation patterns within specific local regions.

## Results

### High-level description of VeloVGI model

Briefly, VeloVGI is a principled variational graph autoencoder(VGAE) based on fine-tuned graph structure to estimate RNA velocity as shown in **Fig 1**. In **Fig 1.a**, this process is designed to handle multiple batches of scRNA-seq data, with batch shapes and cell type colors used to represent the different samples. The method constructs separate inter-batch (in red) and intra-batch (in black) relationships, which are combined to form innovative multi-batch networks. To facilitate analysis, a subset of cells from the network is randomly or node-centricity sampled as input to VeloVGI, while the remaining cells are recovered through subsequent velocity aggregation. In **Fig 1.b**, firstly, for the sampled cells, multiple directed subgraphs are generated using transductive neighbor diffusion strategy. Each directed graph structure corresponds to a mini-batch, with both unspliced and spliced matrices (*u*, s) added as features and jointly passed as inputs to the model. Then, in the specific model, features are extracted in the GCN network to obtain the distribution Z, from which the hidden variable *z* is resampled. The resampled z is assigned to specific induction (green), repression (blue), or steady (pink) states *k* in the spliced-unspliced plane of a particular gene. The decoder estimates the time t, parameter of transcription *α*^(*k*)^, spliced *β*, and degradation *γ* which are calculated jointly (Model specification of Supplementary Methods in VeloVI[22]) to obtain the estimated unspliced and spliced matrices (*ū*(*t*), *s̄*(*t*)), with the similarity of (*ū*(*t*), *s̄*(*t*)) and (*u*, s) as part of the loss target to continuously optimize the parameters. Finally, the velocity is calculated using equation (4). In **Fig 1.c**, the velocity of unsampled cells is recovered by leveraging the known velocity aggregation estimated from the neighborhood of the sampled multi-batch network. In **Fig 1.d**, a variety of biological application interpret model effectiveness such as hierarchical embedding visualization, lineage subcluster, transition probability as well as differential expression.

**Figure 1.**
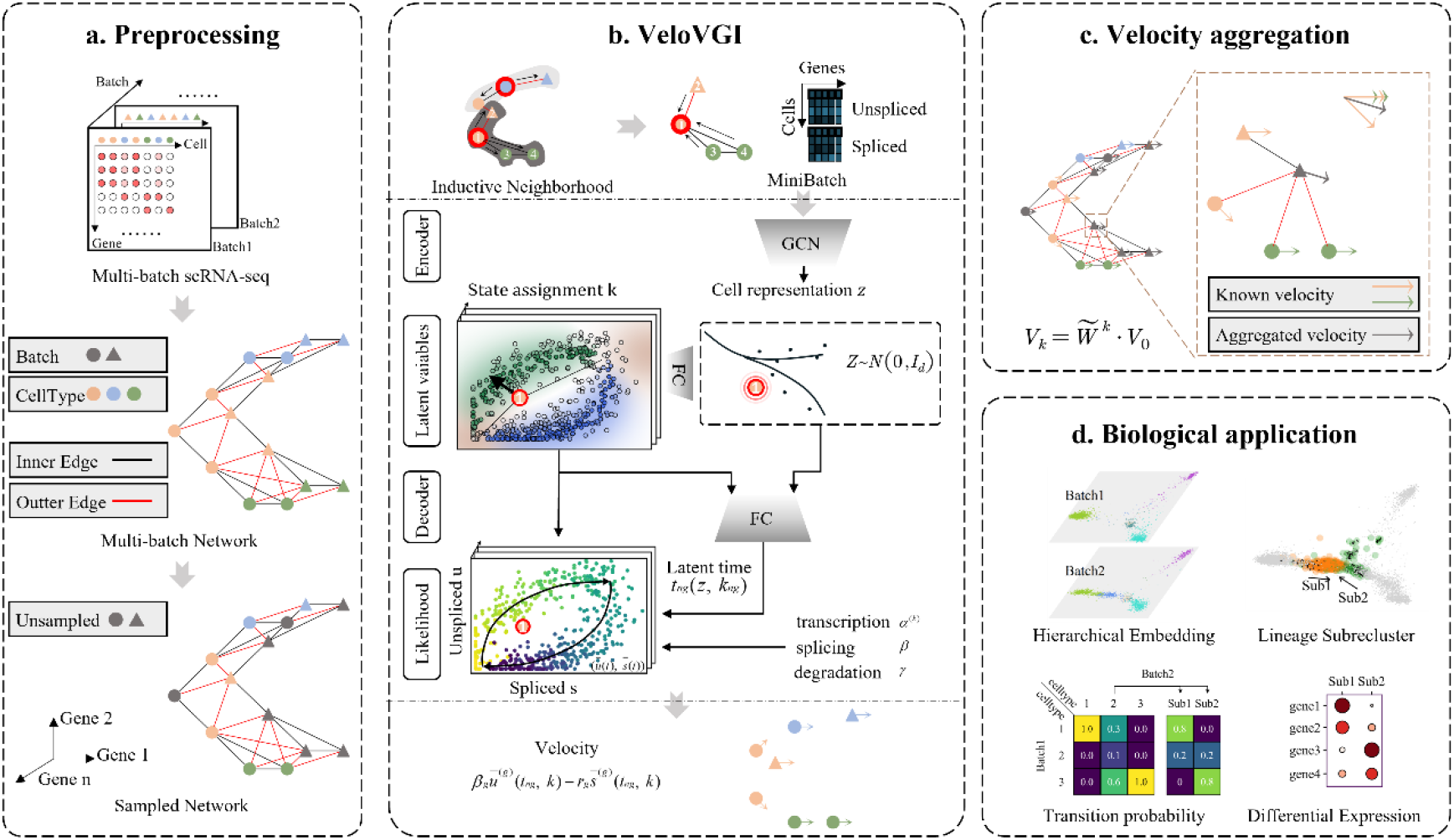
Overview of VeloVGI. a, graph construction of multi-baches network and sampled network in Preprocessing. b, variational graph autoencoder (VGAE) structure and velocity estimation. c, velocity aggregation for unsampled cells. d, a variety of biological application.

### VeloVGI helps to parse the neurodevelopmental heterogeneity of mouse spinal cord tissue across various data sources and injury time points

Firstly, for the spinal cord injury (SCI) dataset, the velocity graph by VeloVGI was applied for analysis, as shown in **Fig 2.a** and **2.b**. These two figures are colored based on cell types and batches, respectively. **Fig 2.c** combines the information from **Fig 2.a** and **2.b**, displaying significant differences in the proportion and distribution of cell types between different batches and highlighting the heterogeneity of cell types before and after injury. Next, the Moscot[24] method was employed to explore the transition relationships of cell types before and after injury, resulting in the transition probability matrix (d). From matrix (d), it can be observed that neural stem cells (NSCs) originate from ependymal cells and astrocyte, corresponding to the sources in the two directions in **Fig 2.a**. Following the concatenation of vector features and coordinate features based on the velocity graph, a clustering was performed once again, yielding lineage-associated subgroups **Fig 2.a** called lineage subcluster. The transition probability matrix **Fig 2.f** demonstrates that the refined NSCs1 and NSCs2 stem respectively from ependymal cells and astrocytes, while **Fig 2.g** indicates the differential expression of marker genes on certain NSCs subtype. NSCs2 exhibit high expression of marker genes in some active neural stem cells (aNSCs), suggesting that these cells are likely aNSCs stimulated post-injury. Additionally, several conclusions aligning with the biological context in **Fig 2.j** were drawn, such as Ependymal cells -> NSCs, TAPs -> Astrocytes, TAPs -> Oligodendrocyte progenitor cells (OPC), TAPs -> Neurons[25–27]. For instances not aligning with the conclusions, this might be due to the non-adjacency of these cells in the dimension reduction graph, which is also one of the key factors influencing the RNA velocity task.

**Figure 2.**
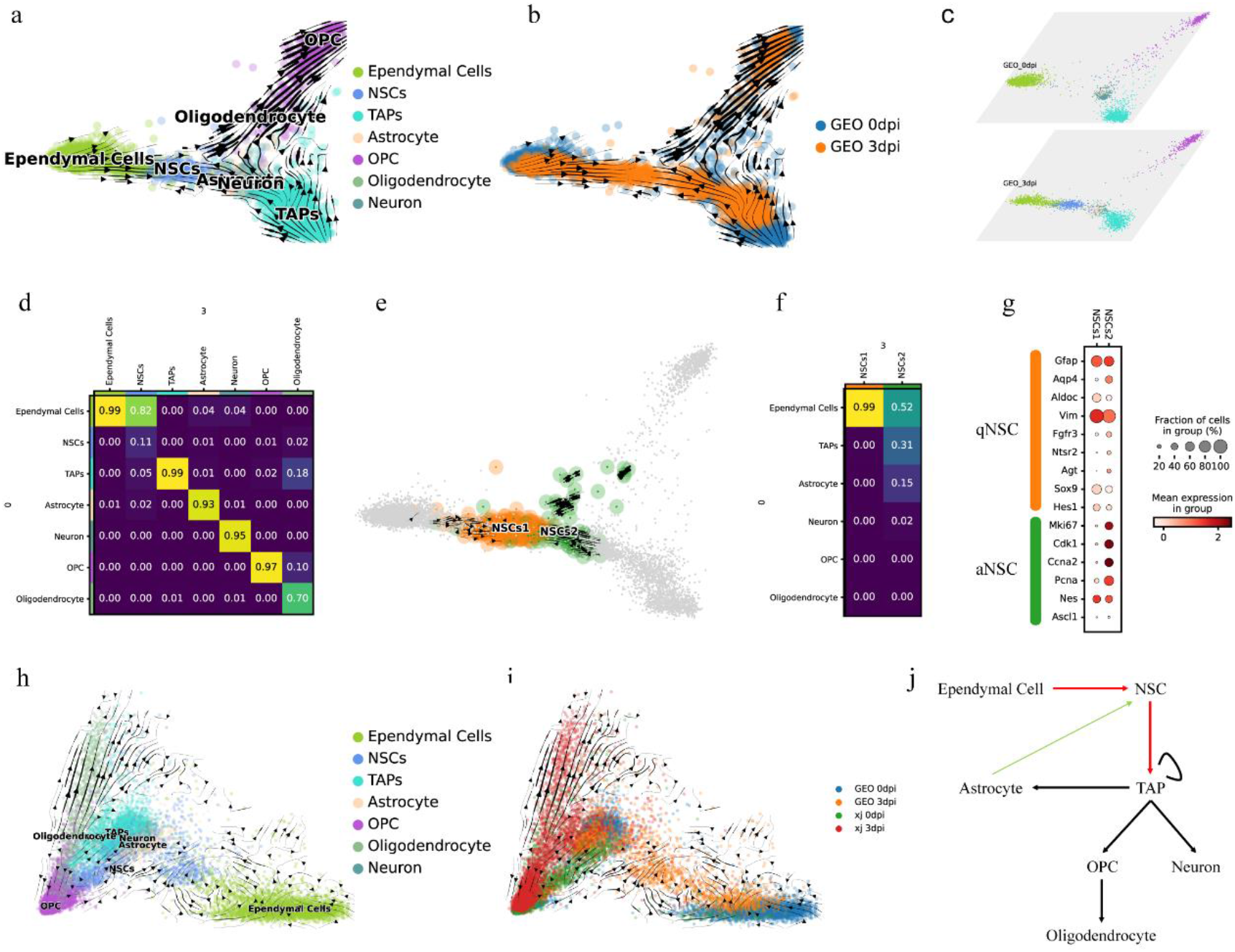
RNA Velocity graph analysis on neural-related cells of spinal cord injury (SCI) tissue with VeloVGI a. and b. show velocity graph with different color to distinguish cell type and batch. c. visualizes the heterogeneity of cell types in different batches by displaying the batches in a hierarchy embedding. d depicts transition probabilities of different cell types across batches from 0 to 3 calculated by Moscot[24]. e. displays velocity graph results of NSCs subtyping based on lineage subcluster (detail in Method) where the transition probabilities and marker gene bubble plot f, g show the clusters difference. h. and i. show velocity graph with additional data processed by the same experimental manipulation. j. illustrate the known difference direction among these related cells.

Furthermore, to gain deeper into the impact of batches on the RNA velocity task and to perform corrections, we conducted identical biological experiments to obtain related data. It can be observed that the data source significantly influences the results, leading to batch effects appearing in the dimension reduction graph. Nevertheless, there still exist instances of cell differentiation that align with the biological context, such as Ependymal cells - NASCs, TAPs -> OPC, OPC -> Oligodendrocyte.

### VeloVGI show the dynamic process of immune related cells during spinal cord injury repair

VeloVGI obtained velocity graph of immune-related cells in mouse spinal cord injury, as shown in **Fig 3.a** and **3.b**. From the perspective of cell types, there is a differentiation trend from Microglia to Macrophage, which aligns with the biological context[28]. Additionally, the batches are displayed here, including un-injury, 12 hours post-injury (pSCI12h), 1 day to 90 days post-injury (pSCI1d∼pSCI90d). Stratifying these cells by time points (**Fig 3.c**) reveals that in the uninjured state, only Microglia cells are present. However, a significant shift occurs after 12 hours of injury stimulation, indicating a trend towards Macrophage differentiation. In the dimensionality reduction plot, cells in the injured state are distanced from the uninjured state and gradually approach it over time, corresponding to the self-repair process after spinal cord injury. Generally, the velocity graph not only illustrates the differentiation process from Microglia to Macrophage cell types but also reflects inter-batch correlations. The neighboring relationships among these batch cells are illustrated in **Fig 3.d**, where connections are established between adjacent batches, and the sample correlations are depicted in **Fig 3.e**.

**Figure 3.**
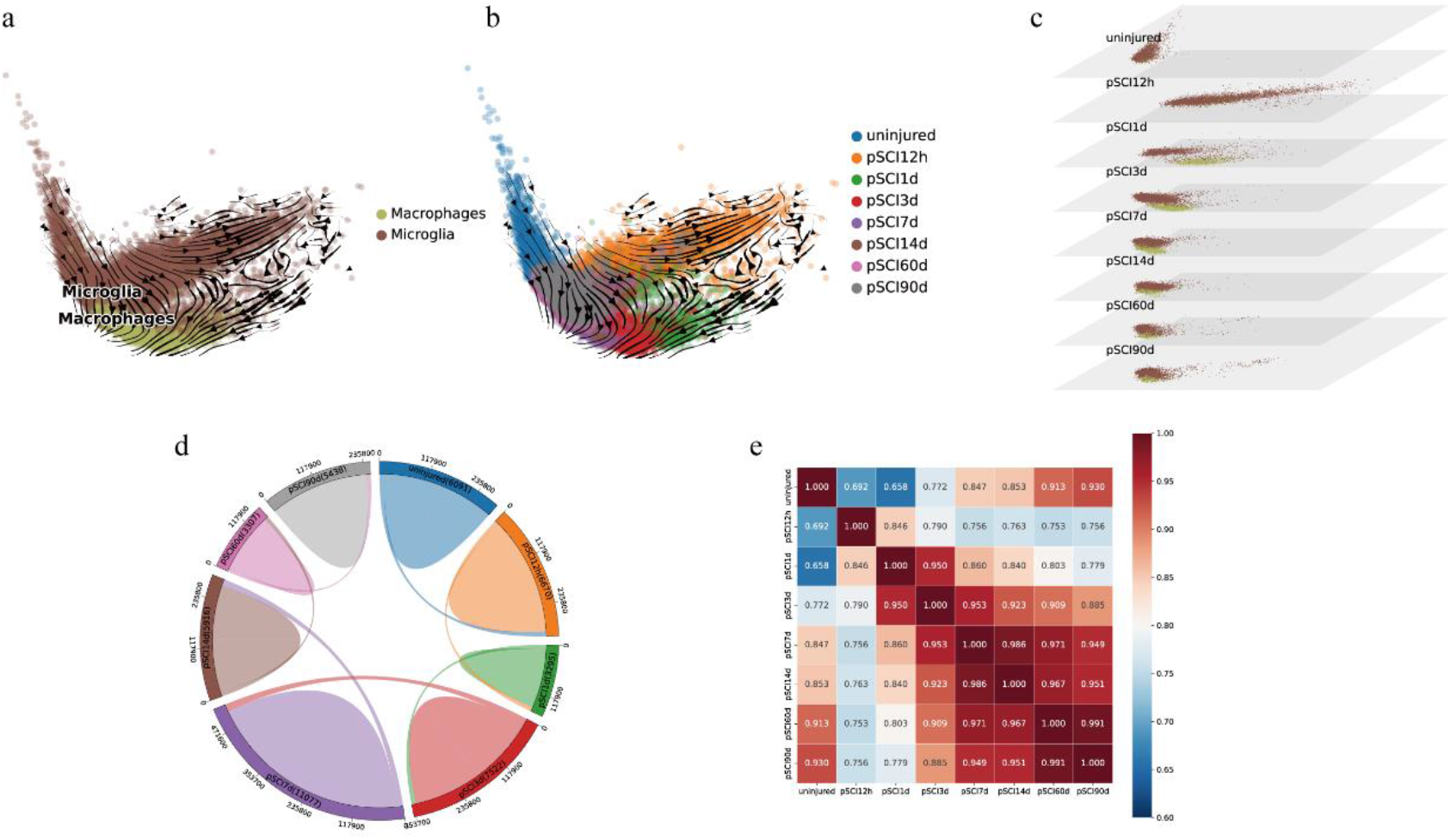
RNA Velocity graph analysis on immune-related cells of spinal cord injury (SCI) tissue with VeloVGI. a. and b. show velocity graph with different color to distinguish cell type and batch. c. visualizes the heterogeneity of cell types in different batches by displaying the batches in a hierarchy embedding. d. depicts the number of neighbors among different batches, establishing batch-to-batch neighbors in chronological order. e. Heatmap illustrating sample correlations between batches.

### VeloVGI reveals the changes in neural system cells during the development of mouse olfactory bulb tissue

The velocity graph of neural system cells in the olfactory bulb tissue generated by VeloVGI is shown in **Fig 4.a** and **4.b**. From a cell type perspective, there is no distinct differentiation relationship between cell types, which is consistent with the biological background of the relevant tissue. Additionally, the figure displays multiple time point batches including embryonic stage (E), 0 day (0d), 2 weeks (2W), and 6 weeks (6W). By grouping cells according to time points (**Fig 4.c**), the gradual developmental changes of the same cell type over time can be observed. The neighboring relationships between these batches are illustrated in **Fig 4.d**, establishing connections between batches from adjacent time points, while the sample correlations are depicted in **Fig 4.e**, where correlations between samples from adjacent time points are stronger.

**Figure 4.**
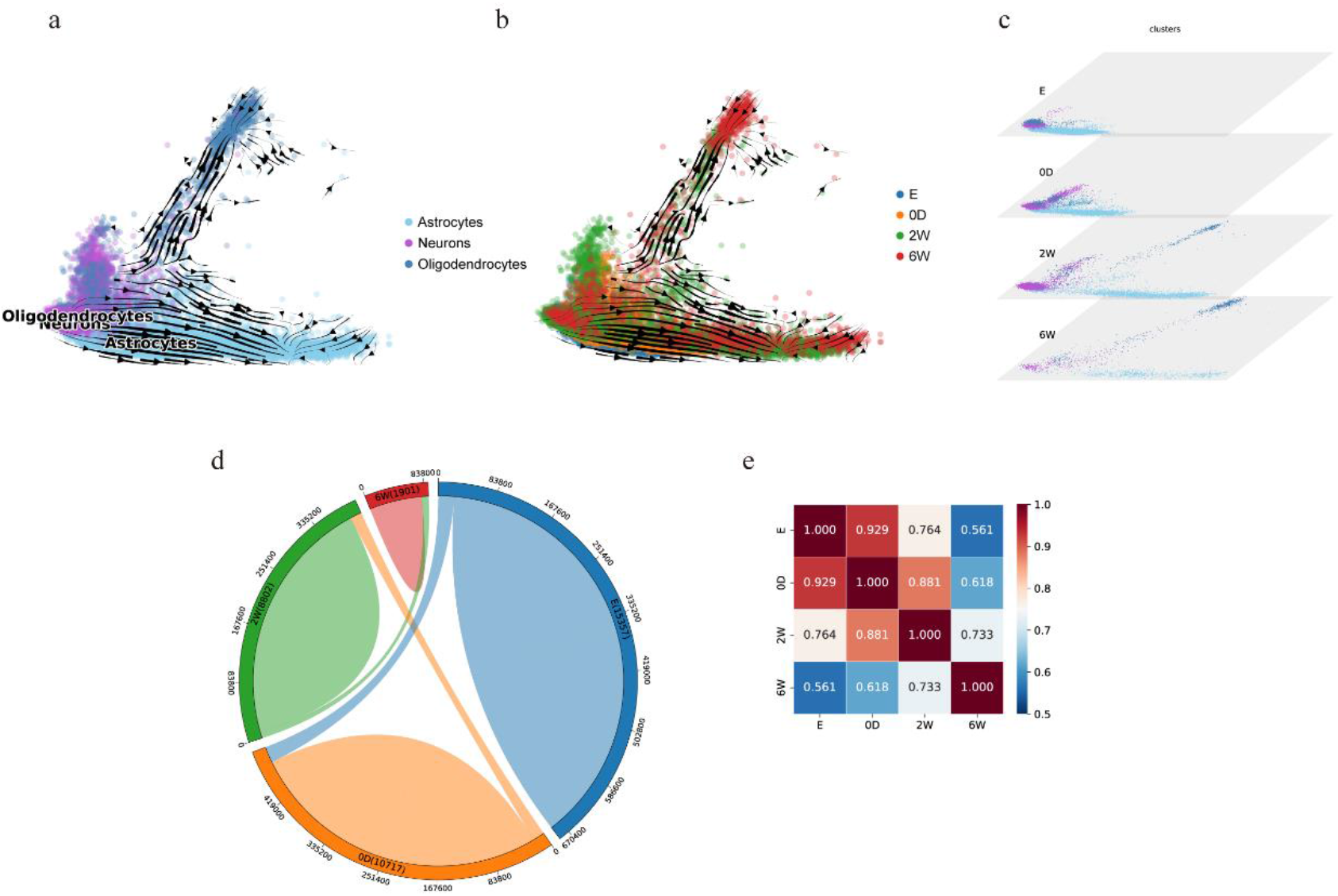
RNA Velocity graph analysis on neural system cells of spinal cord injury (SCI) tissue with VeloVGI. a. and b. show velocity graph with different color to distinguish cell type and batch. c. visualizes the heterogeneity of cell types in different batches by displaying the batches in a hierarchy embedding. d. depicts the number of neighbors among different batches, establishing batch-to-batch neighbors in chronological order. e. Heatmap illustrating sample correlations between batches.

### VeloVGI demonstrates accurate RNA velocity estimation results in diverse data backgrounds

Finally, we apply VeloVGI to datasets from multiple backgrounds, all of which included time series batches. Our method obtained relatively accurate results for these datasets.

On the dentategyrus dataset, although VeloVGI differentiated in opposite directions in microglia and endothelial cells compared to scVelo stochastic mode, there is currently no specific differentiation relationship between the two, which has little impact on the overall accuracy of the RNA velocity results. In addition, VeloVGI can more accurately capture the differentiation direction from OPC (Oligodendrocyte progenitor cell) to OL (Oligodendrocyte) while maintaining the accurate direction from Granule immature to Granule mature, Radial Gila like to Astrocytes (**Fig. 5a**).

**Figure 5.**
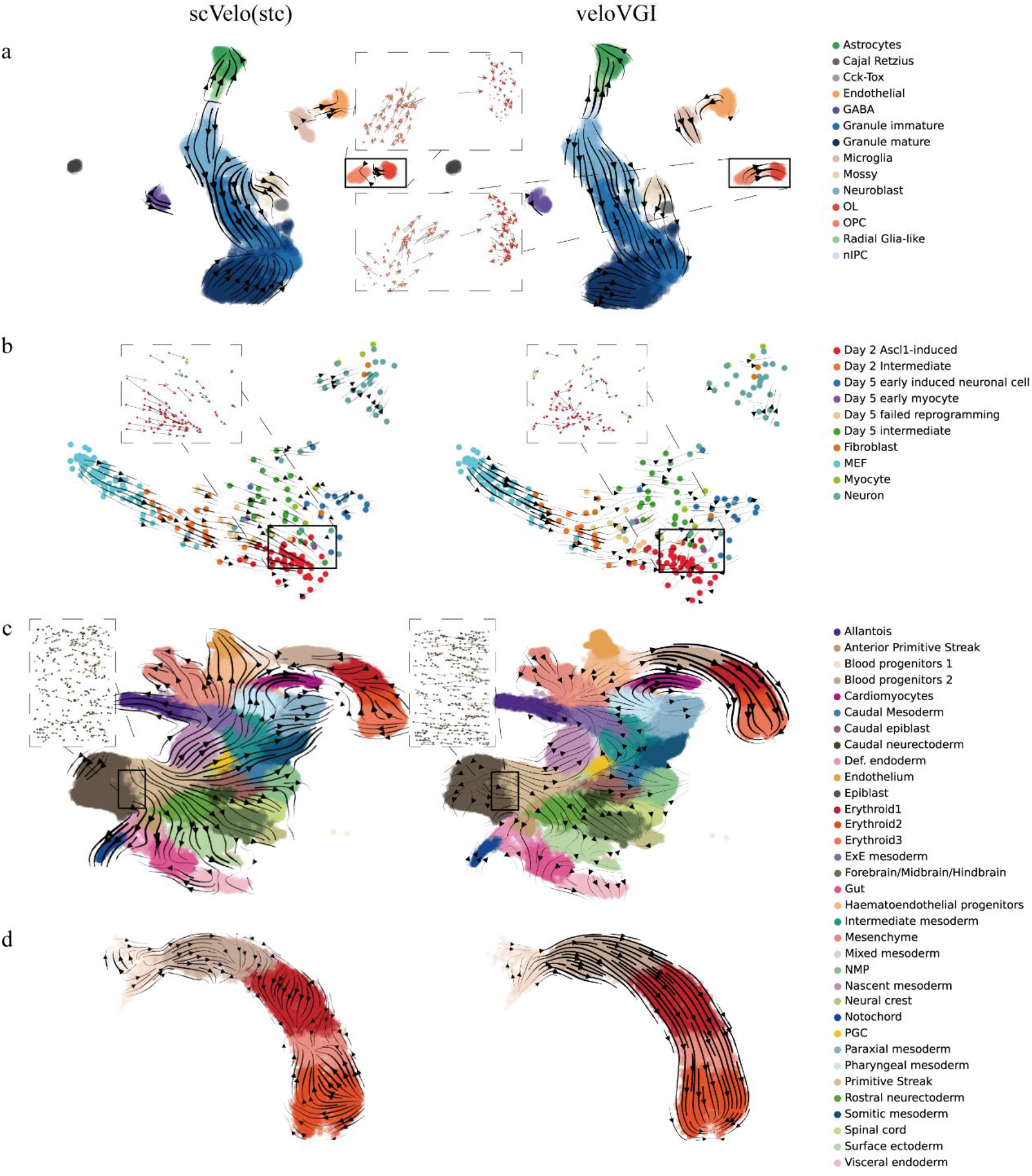
Comparison of scVelo(stc) and VeloVGI for RNA velocity analysis. Analysis on dentategyrus(a), mef reprogramming(b), gastrulation(c), gastrulation erythroid(d). Key parts corrected by VeloVGI are marked with dashed boxes. scVelo(stc) means scVelo with the stochastic.

For the mef reprogramming data, the adjacent time points in the four time point batches of 0, 2, 5, and 22 days are more similar (**Fig. 6a**). For this, we establish neighbors in two adjacent batches (**Fig. 6b, c**), and the neighbor relationship has indicated the approximate direction of differentiation. Although scVelo can obtain a smooth and consistent ground velocity map, for the known differentiation direction “mef → day 2 intermediate → day 2 Ascl1 induced → day 5 intermediate → day 5 early induced neural cells → neuron”, there is a lack of transition from “day 2 Ascl1 induced → day 5 intermediate”, which VeloVGI was able to compensate for (**Fig. 5b**).

**Figure 6.**
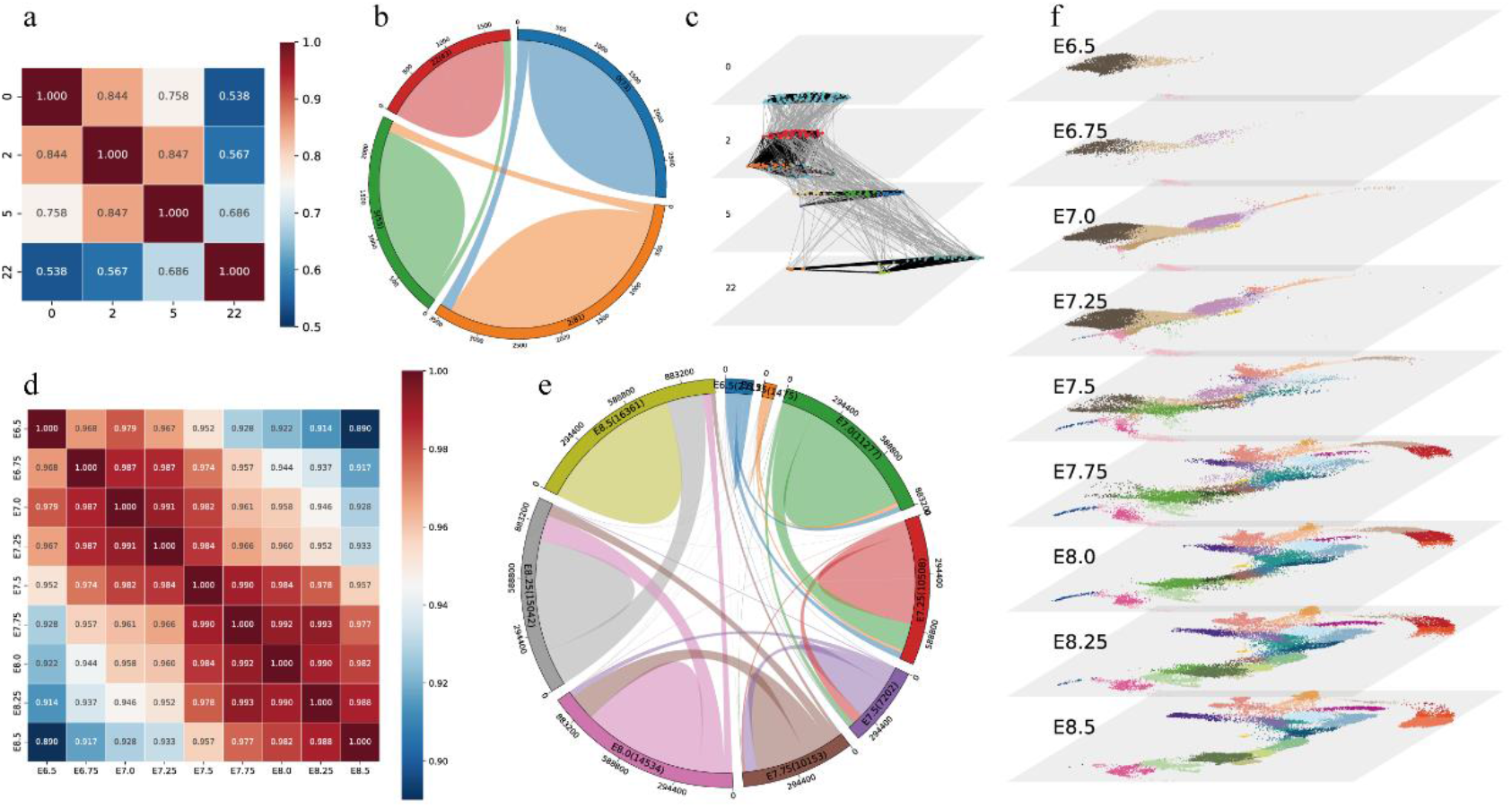
Batch correlation matrix and neighbor edges for mef reprogramming and gastrulation datasets. For mef reprogramming dataset, the adjacent time points in the four time point batches of 0, 2, 5, and 22 days are more similar in correction matrix(a) that neighbor edges are only constructed between them (b, c) Black and gray refer to intra-batch and inter-batch neighbor relationships, respectively. For gastrulation dataset, same as before, with high sample similarity at adjacent time points and overall correlation above 0.89(d), default neighbor construction method of scVelo is used to establish neighbor relationship among all batches (e). What’s more, batch-by-batch UMAP plot demonstrate the dynamics of cell types over time (f).

For the gastrulation data, sampling was conducted every 0.25 days from 6.5 to 8.5 days during the embryonic period. The similarity between the samples was high, with all similarities above 0.89 and the highest similarity between adjacent batches (**Fig. 6d**). Therefore, we do not specify the establishment of neighbors between adjacent batches, but instead use the default neighbor construction method of scVelo to establish neighbor relationship among all batches (**Fig. 6e**). More neighbor relationships can be automatically established between adjacent batches, and the gradual differentiation of cells over time can also be seen from the dimensionality reduction results of batches (**Fig. 6f**). And the velocity graph can more accurately indicate the relationship between cell differentiation (**Fig. 5c**). Compared with scVelo, VeloVGI can capture the transition from cluster Epiblast to Primitive Streak (in the dashed box), while perfectly restoring the erythroid lineage (**Fig. 5d**).

### Comparison experiment

In order to quantitatively compare with other methods in terms of metrics, we introduced CBDir and ICVCoh proposed by the VeloAE[7]. Building upon these metrics, we developed BCBDir and BICVCoh, which are specifically designed to evaluate the impact of RNA velocity on batch datasets. described in the specific formulas in method conducted metric comparisons for all the data presented in the previous sections, as summarized in Tables 1, 2, 3, and 4. In these tables, “N” represent that LatentVelo can’t finish calculation on these datasets for computing power. In Table1,3, “F” represents that there is no known differential direction in these datasets.

**Table 1.** CBDir metric of comparison experiment

| Method\Dataset | sci geo | sci all | sci timeseries | olfactory bulb | erythroid lineage | gastrulation | dentategyrus | mef reprogramming |
| --- | --- | --- | --- | --- | --- | --- | --- | --- |
| scVelo(det) | -0.035 | -0.074 | -0.119 | **F** | -0.028 | -0.280 | 0.301 | 0.313 |
| scVelo(dyn) | -0.009 | -0.061 | -0.132 | **F** | -0.299 | -0.241 | 0.268 | 0.410 |
| scVelo(stc) | -0.020 | -0.070 | -0.141 | **F** | 0.149 | -0.254 | 0.017 | **0.576** |
| VeloAE | -0.070 | -0.030 | -0.043 | **F** | **0.703** | 0.463 | 0.037 | 0.515 |
| DeepVelo | **0.046** | -0.082 | -0.074 | **F** | 0.332 | -0.352 | -0.128 | 0.092 |
| LatentVelo | -0.085 | -0.136 | **N** | **F** | -0.634 | **N** | 0.040 | 0.595 |
| VeloVAE | 0.019 | -0.075 | -0.127 | **F** | -0.248 | -0.211 | 0.131 | 0.558 |
| VeloVI | 0.020 | -0.116 | -0.112 | **F** | 0.060 | -0.495 | 0.408 | 0.063 |
| VeloVGI | 0.031 | **0.058** | **0.128** | **F** | 0.609 | **0.718** | **0.614** | 0.549 |

**Table 2.**
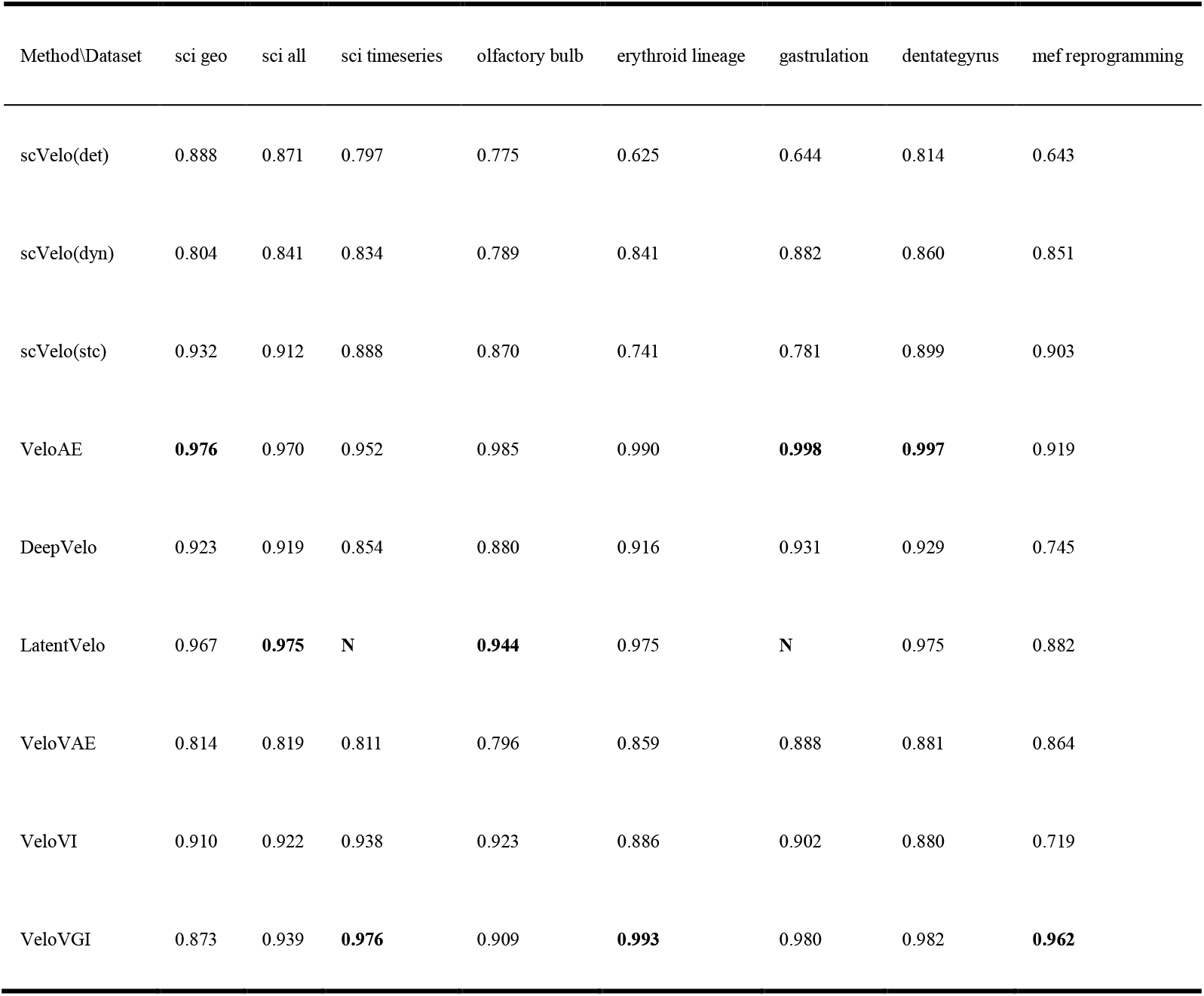
ICVCoh metric of comparison experiment

**Table 3.** BCBDir metric of comparison experiment

| Method\Dataset | sci geo | sci all | sci timeseries | olfactory bulb | erythroid lineage | gastrulation | dentategyrus | mef reprogramming |
| --- | --- | --- | --- | --- | --- | --- | --- | --- |
| scVelo(det) | 0.022 | -0.202 | -0.111 | **F** | -0.028 | -0.281 | 0.233 | 0.322 |
| scVelo(dyn) | -0.021 | 0.127 | -0.138 | **F** | -0.279 | -0.221 | 0.292 | 0.285 |
| scVelo(stc) | 0.024 | -0.251 | -0.128 | **F** | 0.143 | -0.252 | 0.007 | 0.429 |
| VeloAE | -0.088 | **0.251** | -0.060 | **F** | **0.721** | 0.412 | 0.175 | **0.687** |
| DeepVelo | -0.034 | -0.248 | -0.077 | **F** | 0.329 | -0.330 | -0.085 | 0.311 |
| LatentVelo | -0.073 | -0.417 | **N** | **F** | -0.628 | **N** | 0.058 | 0.527 |
| VeloVAE | **0.079** | -0.018 | -0.126 | **F** | -0.224 | -0.200 | 0.287 | 0.417 |
| VeloVI | 0.038 | -0.025 | -0.125 | **F** | 0.063 | -0.487 | 0.489 | -0.014 |
| VeloVGI | 0.047 | 0.030 | **0.097** | **F** | 0.611 | **0.710** | **0.669** | 0.468 |

**Table 4.** BICVCoh metric of comparison experiment

| Method\Dataset | sci geo | sci all | sci timeseries | olfactory bulb | erythroid lineage | gastrulation | dentategyrus | mef reprogramming |
| --- | --- | --- | --- | --- | --- | --- | --- | --- |
| scVelo(det) | 0.689 | 0.602 | 0.792 | 0.753 | 0.591 | 0.618 | 0.783 | 0.274 |
| scVelo(dyn) | 0.645 | 0.627 | 0.779 | 0.776 | 0.836 | 0.873 | 0.840 | 0.882 |
| scVelo(stc) | 0.803 | 0.718 | 0.877 | 0.859 | 0.718 | 0.766 | 0.877 | 0.776 |
| VeloAE | **0.925** | 0.854 | 0.919 | **0.978** | 0.977 | **0.996** | **0.996** | 0.920 |
| DeepVelo | 0.801 | 0.641 | 0.817 | 0.869 | 0.911 | 0.925 | 0.908 | 0.723 |
| LatentVelo | 0.919 | **0.888** | **N** | 0.939 | 0.962 | **N** | 0.966 | 0.631 |
| VeloVAE | 0.654 | 0.576 | 0.743 | 0.783 | 0.854 | 0.879 | 0.845 | **0.926** |
| VeloVI | 0.828 | 0.769 | 0.926 | 0.918 | 0.878 | 0.896 | 0.851 | 0.528 |
| VeloVGI | 0.840 | 0.863 | **0.961** | 0.894 | **0.989** | 0.978 | 0.971 | 0.924 |

**Table 5.** Datasets Overview

| Dataset | # cells | # gene | Known differentiation direction |
| --- | --- | --- | --- |
| Neural-related cells in mouse spinal cord injury (GEO) | 6997 | 31053 | Ependymal Cells → NSCs → TAPs,<br>TAPs → Astrocyte → NSCs,<br>TAPs → OPC → Oligodendrocyte,<br>TAPs → Neuron |
| Neural -related cells in mouse spinal cord injury (GEO+xj) | 20185 | 31053 | Ependymal Cells → NSCs → TAPs,<br>TAPs → Astrocyte → NSCs,<br>TAPs → OPC → Oligodendrocyte,<br>TAPs → Neuron |
| Spinal cord timeseries | 49316 | 42396 | Microglia → Macrophage |
| Olfactory bulb | 36777 | 31053 | None |
| Dentate gyrus | 2930 | 13913 | OPC → OL |
| Mef reprogramming | 252 | 55416 | mef → day 2 intermediate → day 2 Ascl1-induced<br>→ day 5 intermediate → day 5 early induced<br>neuronal cells → neuron |
| Gastrulation | 89267 | 53801 | Epiblast → Primitive Streak, Erythroid1 → 2 → 3 |
| Gastrulation erythroid | 9815 | 53801 | Erythroid1 → 2 → 3 |

In these above table, the metrics are calculated as the aggregated values of all cells in a data corresponding to the metrics, for ICVCoh/BICVCoh metrics it is averaged over all cells, and for CBDir/BCBDir metrics it is averaged within each known cluster pair and then averaged again for the cluster pairs, specific to the inter-distribution metrics box line plots. The cell-level comparisons can be viewed from a box plot perspective such as **Fig 7**, where the performance of the model is better represented. VeloVGI. VeloVGI performs well on various metric on most data.

**Figure 7.**
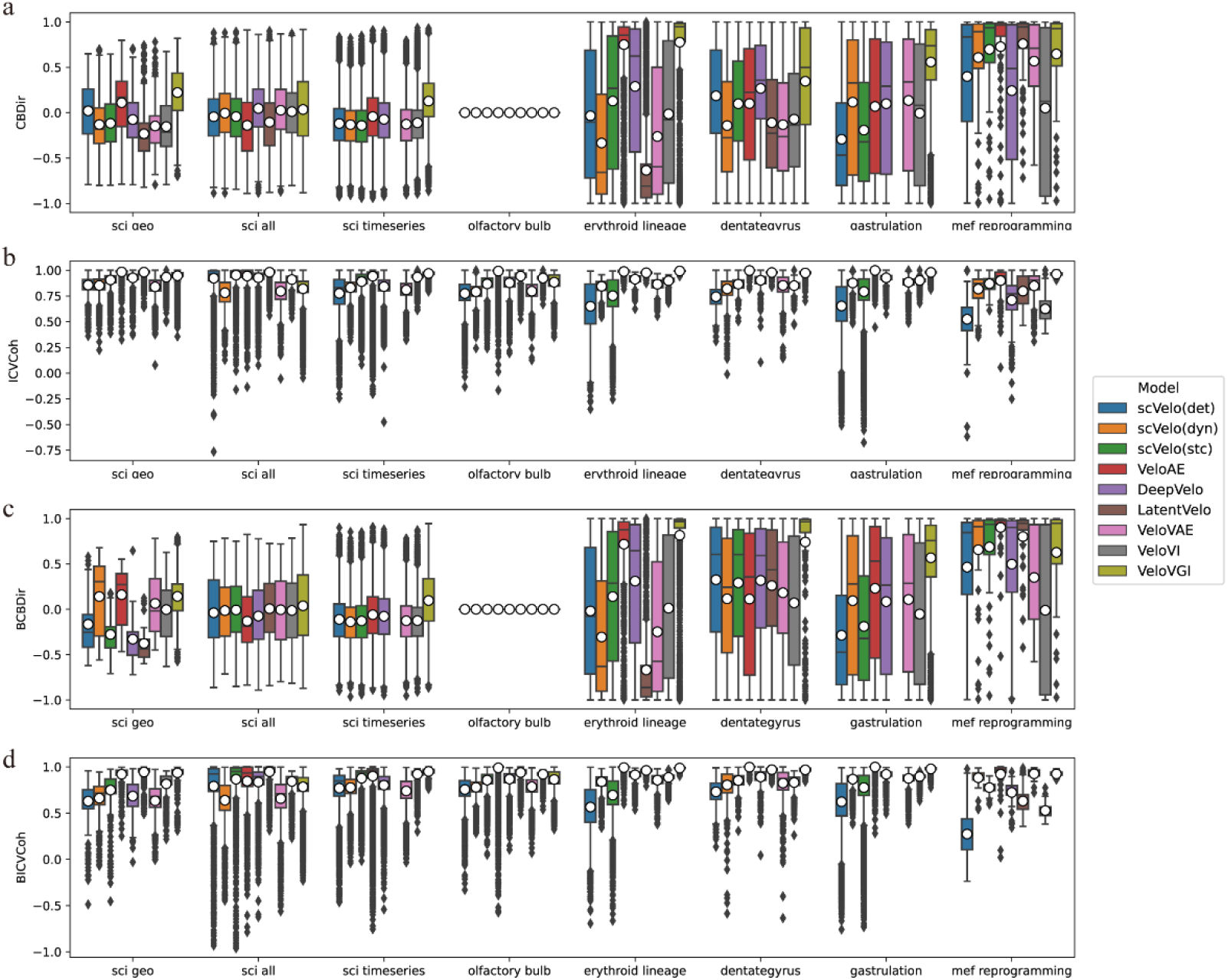
CBDir, ICVCoh BCBDir, BCBDir metric boxplot. Here subplots represent the performance of each of the 4 metrics on each dataset where each subplot has multiple groups of boxplots within it here the groups represent the data, and each group has multiple colors of boxplots within it here the colors represent the RNA velocity model. All metrics range from -1 to 1where the higher the metric the better for the model. Combining all the metrics and data, our model performed best overall.

## Discussion

In this work, we attempt to tackle the challenge of integrating RNA velocity with batch information from the perspective of a neighbor graph structure. We start by constructing a multi-batch network during the preprocessing stage using the mutual nearest neighbors (MNN) technique and the optimal transport theory. In the subsequent parameter estimation stage, we harness the power of graph deep learning techniques. In the subsequent parameter estimation stage, we harness the power of graph deep learning techniques. Finally, we demonstrate the effectiveness of our model through a series of downstream analyses.

In the specific data experiments, we showcase the outcomes of our work. While we demonstrate superior performance compared to existing models on these batch datasets, it’s important to note that the inherent complexity of deep learning models limits our ability to provide in-depth interpretations of the results. The interpretability of deep learning models has been a prominent topic in recent years and remains a focal point for future development. While we modify the CBDir and ICVCoh to BCBDir and BICVCoh, there is a need for further exploration in evaluating the indicators of RNA velocity on batch datasets.

Furthermore, we provide an outlook on the future. Despite the significant advancements made in this work, challenges persist. The limited interpretability of our deep learning model calls for improvements in interpreting model outcomes. Additionally, the applicability of our proposed graph construction strategy under different conditions (such as different time points, treatment operations, etc.) deserves further investigation. We also anticipate extending this graph construction strategy to the integration of single-cell multi-omics data, for instance, employing Weighted Nearest Neighbors (WNN)[29] or inferring RNA velocity in correlated multi-omics data. These directions hold promise for inspiring future research endeavors.

## Materials and Methods

### Data preprocessing

The pre-processing module of all datasets analyzed in this paper does not fully follow the standard procedure of scVelo. Function *scv.pp.filter_and_normalize* with default parameters is used during the procedure of quality control and data transformation. High-quality genes can be reserved that at least 20 cells mRNA expression in both spliced and unspliced count matrix. Principal component analysis could reduce the dimension to accelerate later preprocessing operations.

The crucial step during data preprocessing is neighbor graph construction for batch effect datasets. Unlike traditional graph construction function *scv.pp.neighbors* directly construct KNN ignoring the variations between batches which result in sparse inter-batch neighbor relationships, the innovative graph construction strategy separately construct KNN and mutual nearest neighbors (MNN) among batches, which has been proven to be effective for batch effects[20]. Additionally, employing optimal transportation between relevant batches, followed by MNN, can further enhance the effectiveness of batch effect removal. What’s more, accurately specifying correlation among batches give rise to reliable batch effect removal and RNA velocity estimation. Such as a time series batch dataset can specify the time point sequence during graph construction to capture more reliable neighbor relationships.

For both types of edges, the distance between cells can be calculated using euclidean distance or other suitable metrics. The bidirectional transition probabilities *p_ij_* between cells *i* and *j* can be calculated based on the unidirectional probabilities *p_j|i_* from cell *i* to cell *j* and *p_i| j_* from cell *j* to cell *i*, following equation (1). However, due to the distinct meanings of the two types of edges, their calculation methods also differ. For intra-batch k-nearest neighbors (KNN) construction, the default *sc.pp.neighbor* function provided by scanpy can be utilized with the method described in (2), where *ρ_i_* and *σ_i_* are scaling parameters. For inter-batch mutual nearest neighbors (MNN) construction, the POT package can be used to pass the distance matrix (denoted as ’distance’) between batches a and b, and *ot.emd(Ia, Ib, distances)* can be called to obtain the optimal transfer probabilities matrix. For this matrix, we retain the top K values that represent the highest mutual transition probabilities between each pair of cells.

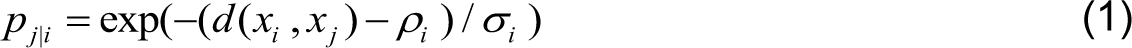

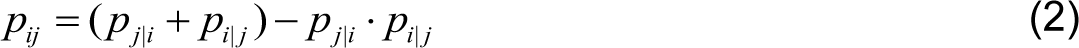

### Parameter inference of VeloVGI

Several models based on the Variational Autoencoder (VAE) framework have been proposed for RNA velocity analysis in single-cell omics data. Notable examples include VeloVAE[10], Pyro-Velocity[11], LatentVelo[12] and VeloVI [22]. The VAE model have demonstrated its effectiveness in removing batch effects [21], making it valuable tools for analyzing scRNA-seq data. In this study, estimate RNA velocity parameters are estimated by VeloVI due to its efficient implementation in scvi-tools [30], a Python library designed for probabilistic analysis of single-cell omics data which is easy to deploy and redevelopment.

The specific estimation process of VeloVI is centered around VAE. The concatenated matrix (*U*, *S*), formed by combining the spliced matrix *S* and the unspliced matrix *U*, serves as the input for the model. The encoder fits the distribution of features and resamples to obtain the hidden layer representation of cells. The decoder first uses this embedding to fit the parameters of the transcription rate and the state of the cell, and then uses the basic RNA velocity assumptions and the derived formulas for cell and gene specificity to fit the estimated (*U^*, *S^*). Finally, model trained by minimizing MSE (Mean Square Error) through gradient descent.

The transcription *α*, splicing *β* and degradation *γ* parameters conform to the differential equations of a kinetic model. These parameters are estimated by fitting the decoder of a Variational Autoencoder (VAE) as an auxiliary task, and the velocity are computed using the estimated parameters by formula (4).

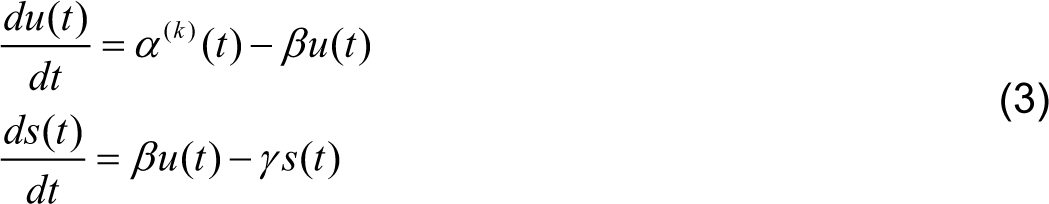

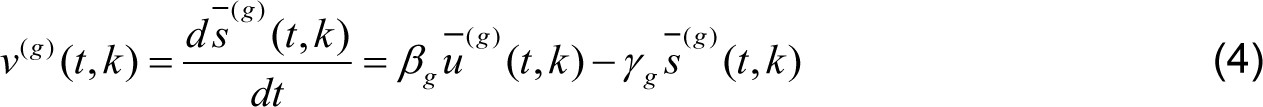

VeloVGI adds a graph representation learning strategy in the encoder section based on it, fully utilizing neighbor relationships to enhance the model’s representation ability, as described in the next section.

### Graph Learning Representation

#### 1. GCN

Graph Convolution Network (GCN)[31] is primarily designed to extract latent representation of nodes by combining original feature of nodes and neighbor relationships, which can be conducted in matrix form as

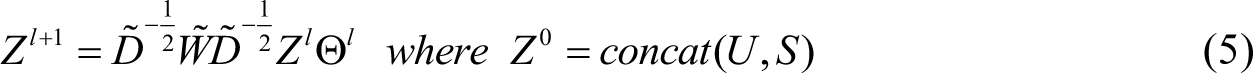

The formula consists of message generation and message aggregation. Message generation is expressed as where *Z ^l^* and Θ*^l^* correspond to feature and weight in *l* -th layer of model. Message aggregation is represented as 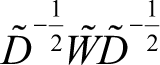 where 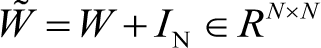 based weighted adjacency matrix additionally adds self-loops and *D̃* is the degree matrix of *W̃*. The formula as a whole implements the feature transformation from layer *l* to layer *l* +1 while *Z*^0^ matrix is the concatenation of unsplice*W* d count matrix *U* and spliced count matrix S.

#### 2. MiniBatch, GraphSAGE

Since scRNA-seq datasets from tissue samples are usually composed of multiple batches and the number of cells may be in the tens or even hundreds of thousands, using GPU for training becomes extremely resource-intensive if all cell features and neighbor relationship graphs are input into the model simultaneously. There for, a minibatch strategy for graph is needed to split whole a graph into many subgraphs to train model. Simple random sampled minibatch is not suitable for this task, which results in the loss of large number of neighbor relationships while every minibatch should preserve sufficient neighbor relationship for model training. After trying many other strategies, an inductive representation learning and neighborhood sampling method called GraphSAGE[23] was found to be well-suited for this task. This strategy, after sampling randomly selected nodes, extends to their surrounding neighbors and generates a directed graph for subsequent message aggregation. The number of neighbors for each GCN layer and the GCN layer count can be adjusted as needed.

The implementation of this graph learning representation part is mainly based on PyTorch Geometric[32].

### Sample and recovery strategy

#### 1. Sample

The presence of a large number of cells is a notable characteristic of multi-batch datasets. In addition to the previous mentioned minibatch strategy to reduce resource pressure in feature extraction, down-sampling during preprocessing can also effectively compress the dataset size and reduce resource consumption. Regarding the specific sampling method, we discovered that random sampling is straightforward and effective after experimenting with various methods, including node centrality sampling.

#### 2. Recovery

For cells that have not been sampled, their velocity-related properties (including the velocity vectors in both high and low dimensions, the relevant parameters in the underlying assumptions, etc.) can be can be obtained by calculating the average of the corresponding properties of the sampled neighboring cells. This process is similar to the moment operation in the scVelo package preprocessing process.

### Lineage subcluster

The typical subcluster based on gene expression values often involves subjective decisions when determining whether further sub-clustering is necessary. It requires choices of resolution and cluster count without clear reference points. In this regard, lineage subcluster offer a solution to these challenges. When accurate velocity graphs are available, some cellular clusters may clearly exhibit multiple distinct differentiation trends. For example, as seen in **Fig 2a**, NSCs exhibit two distinct subgroups. This observation indicates the presence of various differentiation potentials within these clusters, a phenomenon we term lineage re-cluster. These lineage subcluster are defined based on the differentiation direction presented in the lineage or developmental trajectory.

The method for identifying lineage subgroups is straightforward. Firstly, it is necessary to determine which major clusters have the potential for lineage subgroups. This assessment can be made by observing the velocity graph to determine whether a particular cluster exhibits consistent and gradual changes in velocity vectors. In this context, “consistency” can be inferred by examining whether the in-cluster coherence (ICVCoh) metric of the major cluster is sufficiently large. The concept of “gradual changes” can be evaluated by measuring the similarity between the velocity vectors of all cells within the major cluster and the average velocity vector of the cluster.

Next, the specific segmentation of lineage subgroups needs to be established. In this context, we perform conduct clustering on concatenated features X_lineage_subrecluster_ = [*V_embedding_*, *X_embedding_*]. The optimal number of clusters for lineage subgroups is determined by selecting the clustering solution with the highest silhouette coefficient. This process enhances the accuracy of identifying the structure of lineage subgroups.

### Evaluation Metric

The differentiation relationship in the biological background, which play a significant role in RNA velocity result assessment, can evaluate the validity of RNA velocity methods on a reduced dimensional visualization graph such as UMAP or tSNE. Furthermore, cross-boundary direction correctness (CBDir) and in-cluster coherence (ICVCoh) are currently recognized as a relatively reliable evaluation metrics that transform fuzzy visualizations into precise values[7, 8].To elaborate, CBDir measures cosine similarity of velocity and expression difference among neighbor cells on the ideal cluster differentiation boundary and specific direction within the heterogeneous cluster to evaluate how likely a cell can develop towards other neighbor target cells. ICVCoh, on the other hand, assesses velocity consistency among neighbor cells within the homogeneous cluster, representing the smoothness of cell velocity.

Boundary cells should be defined before computing CBDir, which requires pairs of cell clusters as input of ground truth development directions. Boundary cells from A to B represent the set C*_A → B_* as

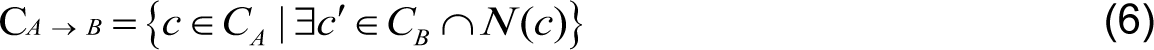

The formula for computing CBDir score is

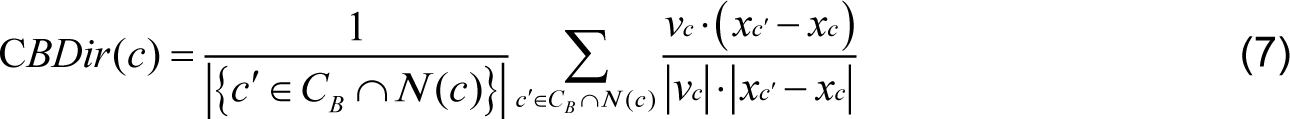

The formula for computing ICVCoh score is

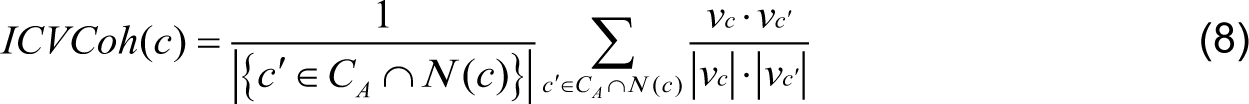

In the above formula, *N*(*c*) is a set of neighbors of cell *c*. *xc*′ and *xc* represent the expression of source cell *c* and target cell *c*′ in low-dimensional space in UMAP, tSNE or other embedding space.

However, in the real batch datasets, since the inter-batch neighbor relationships are much fewer than the intra-batch neighbor relationships, the two existing metrics may focus too much on intra-batch cell velocity performance and ignore the velocity results of batch effect integration. To address this issue, we improved on the two previous metrics by constructing BCBDir and BICVCoh to care only about inter-batch neighbor relationships and provide a more effectively assessment of the performance of RNA velocity on datasets with batch effects. The slight modification is that target cell *c*′ should not belong to the same batch with source cell *c*. The new boundary cell set 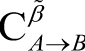, BCBDir, BICVCoh are as follow:

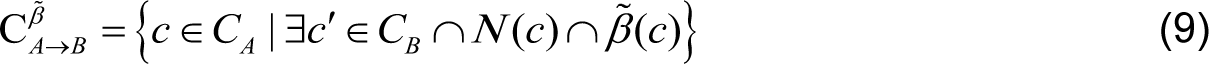

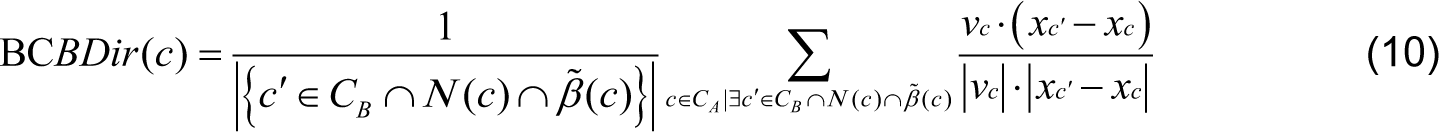

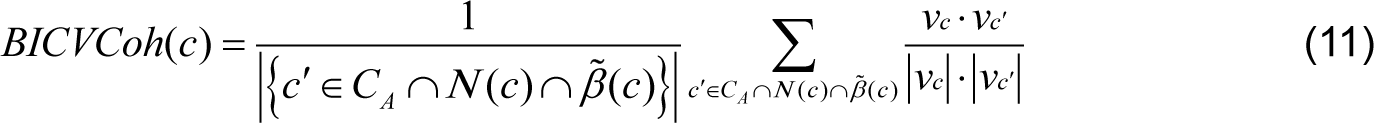

## Datasets

To evaluate our methods, scRNA-seq datasets with multiple batches from different biological systems are collected and input into our model enrolled: spinal cord. These datasets contain known differentiation orders for all or part of the cell clusters that we can validate the result correctness of estimated velocity by visualization plot and metric value.

The neural-related cells dataset is from two source, GEO and our experiment (label as xj). The GEO dataset is accessible from GSE162610 with 10X genomics platform. The xj dataset sequenced on BD Rhapsody platform will be released in the near future. Both of them are from mouse spinal cord injury tissue with many cell types while only neural-related cells are select to detect the latent differential relationship.

The spinal cord timeseries dataset is downloaded from GSE189070 as 10X genomics platform fastq file. The immune-related cells are select to detect the latent differential relationship.

The olfactory bulb dataset will be released in the near future sequenced on BD platform. The neural-related cells are select to detect the latent differential relationship.

The above datasets need to be processed upstream to get spliced/unspliced expression matrix from fastq files, where velocyto[3] is used after cellranger 6.1.2 for 10X genomics platform or rhapsody 1.10 for BD Rhapsody platform. The following datasets can be accessible as spliced/unspliced expression matrix format directly.

The dentate gyrus dataset is one of the most classical datasets for RNA velocity task that can directly downloaded from scVelo packages with scvelo.datasets.dentategyrus. The transition from OPC (Oligodendrocyte progenitor cell) to OL(Oligodendrocyte) is the known differentiation direction that accurately estimated by our method.

The mef reprogramming dataset is the result of one of the first methods for designing time-series RNA-seq experiments and preprocessed[13].The time series of the dataset is 0, 2, 5, 22 days after reprogramming. The known transition is from mouse embryonic fibroblasts (MEF) to neuronal cells that expressed as “mef →day 2 intermediate → day 2 Ascl1-induced → day 5 intermediate → day 5 early induced neuronal cells → neuron”.

The mouse gastrulation and gastrulation erythroid dataset are both from scVelo package, where the batch interval is 0.25 day which is so small that the K nearest neighbors for all total dataset is more suitable for better result.

## Code Availability

VeloVGI is an open-source Python package available at GitHub, https://github.com/HuangDDU/velovgi_workstation. All the analysis notebooks for reproducing the results are also available in this repository.

## Hardware configuration

This work’s hardware configuration is supported by high-performance computing platform of Xidian University with GPU of A100 and V100s.

## Author Contributions

All authors contributed to the article. LY and CZ initiated and envisioned the study. ZH, XG and LY formulated the model. ZH was responsible for implementing the algorithm and collecting dataset. CZ and JQ conceived and carried out the biological experiments. ZH and LY were responsible for writing the manuscript, which was subsequently reviewed, edited, and approved by all authors.

## Classification

Computer Sciences; Cell Biology

## Acknowledgments

Thanks to all those who maintain excellent databases and to all experimentalists who enabled this work by making their data publicly available.

---

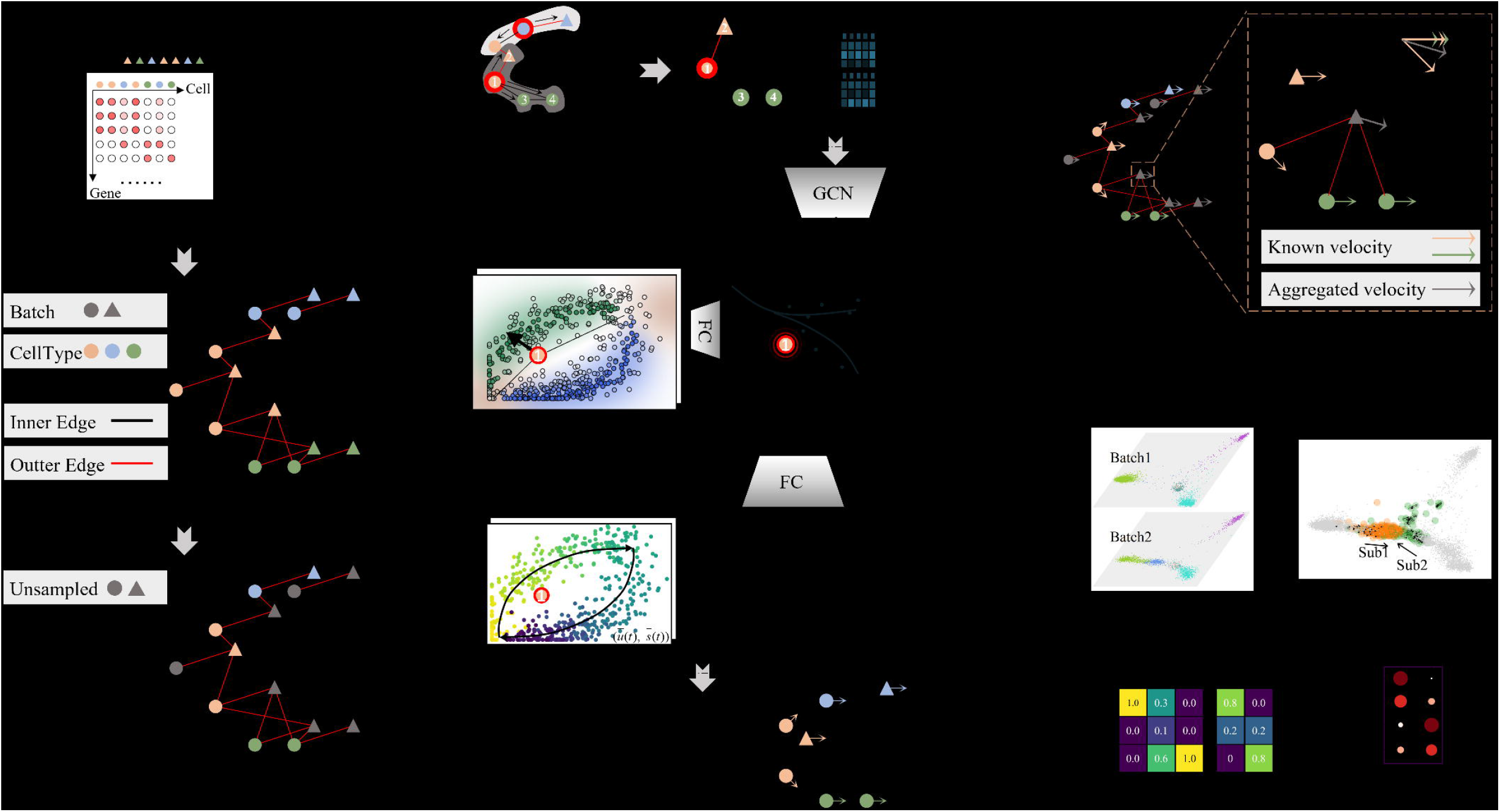

## References

1. J. A. Griffiths, A. Scialdone, J. C. Marioni, Using single-cell genomics to understand developmental processes and cell fate decisions. Molecular systems biology 14, e8046 (2018).

2. W. Saelens, R. Cannoodt, H. Todorov, Y. Saeys, A comparison of single-cell trajectory inference methods. Nature biotechnology 37, 547–554 (2019).

3. G. La Manno et al., RNA velocity of single cells. Nature 560, 494–498 (2018).

4. V. Bergen, M. Lange, S. Peidli, F. A. Wolf, F. J. Theis, Generalizing RNA velocity to transient cell states through dynamical modeling. Nature biotechnology 38, 1408–1414 (2020).

5. M. Lange et al., CellRank for directed single-cell fate mapping. Nature methods 19, 159–170 (2022).

6. X. Qiu et al., Mapping transcriptomic vector fields of single cells. Cell 185, 690–711. e645 (2022).

7. C. Qiao, Y. Huang, Representation learning of RNA velocity reveals robust cell transitions. Proceedings of the National Academy of Sciences 118, e2105859118 (2021).

8. M. Gao, C. Qiao, Y. Huang, UniTVelo: temporally unified RNA velocity reinforces single-cell trajectory inference. Nature Communications 13, 6586 (2022).

9. Z. Chen, W. C. King, A. Hwang, M. Gerstein, J. Zhang, DeepVelo: Single-cell transcriptomic deep velocity field learning with neural ordinary differential equations. Science Advances 8, eabq3745 (2022).

10. Y. Gu, D. Blaauw, J. D. Welch, Bayesian inference of RNA velocity from multi-lineage single-cell data. bioRxiv, 2022.2007. 2008.499381 (2022).

11. Q. Qin, E. Bingham, G. La Manno, D. M. Langenau, L. Pinello, Pyro-Velocity: Probabilistic RNA Velocity inference from single-cell data. bioRxiv, 2022.2009. 2012.507691 (2022).

12. S. Farrell, M. Mani, S. Goyal, Inferring single-cell dynamics with structured dynamical representations of RNA velocity. bioRxiv, 2022.2008. 2022.504858 (2022).

13. J. Ding, N. Sharon, Z. Bar-Joseph, Temporal modelling using single-cell transcriptomics. Nature Reviews Genetics 23, 355–368 (2022).

14. M. D. Luecken et al., Benchmarking atlas-level data integration in single-cell genomics. Nature methods 19, 41–50 (2022).

15. D. He et al., Alevin-fry unlocks rapid, accurate and memory-frugal quantification of single-cell RNA-seq data. Nature Methods 19, 316–322 (2022).

16. P. Melsted et al., Modular, efficient and constant-memory single-cell RNA-seq preprocessing. Nature biotechnology 39, 813–818 (2021).

17. V. Bergen, R. A. Soldatov, P. V. Kharchenko, F. J. Theis, RNA velocity—current challenges and future perspectives. Molecular systems biology 17, e10282 (2021).

18. C. Soneson, A. Srivastava, R. Patro, M. B. Stadler, Preprocessing choices affect RNA velocity results for droplet scRNA-seq data. PLoS computational biology 17, e1008585 (2021).

19. G. Schiebinger et al., Optimal-transport analysis of single-cell gene expression identifies developmental trajectories in reprogramming. Cell 176, 928–943. e922 (2019).

20. L. Haghverdi, A. T. Lun, M. D. Morgan, J. C. Marioni, Batch effects in single-cell RNA-sequencing data are corrected by matching mutual nearest neighbors. Nature biotechnology 36, 421–427 (2018).

21. R. Lopez, J. Regier, M. B. Cole, M. I. Jordan, N. Yosef, Deep generative modeling for single-cell transcriptomics. Nature methods 15, 1053–1058 (2018).

22. A. Gayoso et al., Deep generative modeling of transcriptional dynamics for RNA velocity analysis in single cells. bioRxiv, 2022.2008. 2012.503709 (2022).

23. W. Hamilton, Z. Ying, J. Leskovec, Inductive representation learning on large graphs. Advances in neural information processing systems 30 (2017).

24. D. Klein et al., Mapping cells through time and space with moscot. bioRxiv, 2023.2005. 2011.540374 (2023).

25. R. Chevreau et al., RNA profiling of mouse ependymal cells after spinal cord injury identifies the oncostatin pathway as a potential key regulator of spinal cord stem cell fate. Cells 10, 3332 (2021).

26. C. Li et al., Temporal and spatial cellular and molecular pathological alterations with single-cell resolution in the adult spinal cord after injury. Signal transduction and targeted therapy 7, 65 (2022).

27. K. Obernier, A. Alvarez-Buylla, Neural stem cells: origin, heterogeneity and regulation in the adult mammalian brain. Development 146, dev156059 (2019).

28. S. David, A. Kroner, Repertoire of microglial and macrophage responses after spinal cord injury. Nature Reviews Neuroscience 12, 388–399 (2011).

29. Y. Hao et al., Integrated analysis of multimodal single-cell data. Cell 184, 3573–3587. e3529 (2021).

30. A. Gayoso et al., A Python library for probabilistic analysis of single-cell omics data. Nature biotechnology 40, 163–166 (2022).

31. T. N. Kipf, M. Welling, Semi-supervised classification with graph convolutional networks. arXiv preprint arXiv:1609.02907 (2016).

32. M. Fey, J. E. Lenssen, Fast graph representation learning with PyTorch Geometric. arXiv preprint arXiv:1903.02428 (2019).

